# Selection of Escherichia coli porin and LPS mutants under exposure to phage T4 facilitates the emergence of β-lactam resistance

**DOI:** 10.1101/2025.10.30.685503

**Authors:** Justine Le-Boulch, Erika Charneau, Anne Chevallereau, Arnaud Gutierrez

## Abstract

Understanding all sources of selective pressure that contribute to the emergence of antibiotic resistance is essential for developing sustainable antimicrobial strategies. Here, we investigated the interaction between phage T4 and Escherichia coli MG1655 to determine whether mutations conferring phage resistance also shape the genetic background for β-lactam resistance. Using experimental evolution, whole-genome sequencing, and targeted genetic reconstructions, we identified mutations in porins and lipopolysaccharide (LPS) biosynthesis as the predominant routes to phage T4 resistance. Precise allelic replacements and isogenic strain comparisons demonstrated that these mutations not only protect against phage predation but also create a genetic context that facilitates the emergence of β-lactam resistance, including resistance to carbapenems. Together, these findings provide compelling evidence that phage-driven selection can establish bacterial genetic backgrounds predisposed to antibiotic resistance. This work highlights the evolutionary risks associated with phage therapy and underscores the need to account for genetic trade-offs when developing alternative antimicrobial strategies.

## Introduction

The interaction between bacteria and their viruses, bacteriophages (phages), is recognized as a major force shaping microbial ecosystems and significantly influencing bacterial evolution [1,2]. Depending on the environment, phages are at least as numerous as bacteria, and some estimates suggest that the global number of phages is around 10^31^, ten times higher than that of bacteria, making them the most abundant biological entities on Earth [3]. As one of the primary natural predators of bacteria, phages exert strong selective pressure on bacterial populations [4,5]. Phage predation is believed to account for 20% to 40% of daily bacterial lysis, depending on environmental conditions [6]. The evolutionary dynamics of phage-bacteria interactions are often characterized as antagonistic coevolution, particularly in the form of an evolutionary arms race. In response to phage predation, bacteria evolve mechanisms to evade infection, while phages, in turn, adapt to overcome bacterial defenses. This ongoing interplay affects not only bacterial resistance to phages but also shapes broader evolutionary trajectories through mechanisms such as second-order selection, pleiotropy, and epistasis—ultimately giving rise to traits that influence bacterial adaptability and survival in diverse environments [7,8]. As such, antagonistic coevolution between phages and bacteria represents a major evolutionary force in microbial communities and has the potential to drive the emergence of novel bacterial phenotypes. A critical step in phage-bacteria interactions is the attachment of phages to bacterial surface structures - or surface antigens - such as outer membrane proteins, porins, lipopolysaccharides (LPS), and capsules [9]. While these structures serve as phage receptors, they are also essential for the interactions between bacteria and their environment. In Gram-negative bacteria, porins are among the primary constituents of the outer membrane [10]. These proteins form channels that facilitate the uptake of small molecules, including nutrients and essential ions. Porins can also act as receptors for specific phages, providing an anchor point for phage to inject genetic material [9]. This dual function, as both nutrient transporters and phage receptors, makes porins a key element in phage-bacteria evolutionary dynamics. Moreover, certain antibiotics, such as β-lactams or fluoroquinolones, are described to use porins as entry pathways to access the bacterial cell and inhibit essential functions [11,12]. As a result, selection for porin modifications or loss to avoid phage attachment may also coincidentally reduce antibiotic permeability, thereby altering the bacterial antibiotic resistance profile.

One of the most critical consequences of bacterial adaptability, especially concerning human health, is the emergence and spread of antibiotic resistance [13]. Bacteria have evolved numerous mechanisms that contribute to multidrug resistance, posing serious public health concerns [14]. In Gram-negative bacteria, a key resistance mechanism involves reduced antibiotic uptake due to porin loss or structural alteration [15]. When combined with β-lactamase expression, decreased permeability to β-lactams can result in high levels of resistance [16]. Although carbapenems are not efficiently hydrolysed by β-lactamase aside carbapenemase such as NDM-1 or KPC2, their expression with reduced outer membrane permeability can lead to clinically significant resistance and treatment failure [16]. For instance, the co-expression of OXA-48 family β-lactamases and altered porin profiles was reported to confer resistance to carbapenem antibiotics such as meropenem and ertapenem [17,18]. Selection of carbapenem resistance caused by loss of porins and expression of AmpC β- lactamase was reported after ertapenem treatment [19]. This phenotype more commonly referred as “Non-carbapenemase carbapenem resistance” (NC-CR) was also described to be mediated by selection of ceftazidime resistance in a *Klebsiella* clinical isolate [20]. Even though NC-CR was linked to treatment failure, in *Enterobacteriaceae* NC-CR isolates are primarily associated with asymptomatic carriage [21] and are less virulent than carbapenemase-producing strains [22].

Given that antibiotic resistance represents a major threat to our medical practice, there is an urgent need for the discovery of new antimicrobial compounds [23], revised antibiotic usage policies [14], and the development of alternative therapies [24]. One such alternative that has regained interest is phage therapy [25], particularly for treating infections such as urinary tract infections and others caused by multidrug-resistant bacteria. Phage therapy offers several advantages are highly specific to target bacteria, effective against antibiotic-resistant strains, and can work synergistically with antibiotics [26]. Additionally, some phages possess biofilm-degrading activity [27], and phage production is likely to be cost-effective [28]. The effectiveness of phage therapy may be compromised by evolution of bacterial resistance to phages, which often occurs through the selection of receptor variants. However, little is known about the consequences of intensifying selective pressure on bacterial surface antigens [29]. In addition, although resistance to phages may limit the ability to eliminate the targeted population, there is increasing interest in leveraging the fitness costs associated with phage resistance [30,31]. This concept, known as phage steering, involves using virulent phages to drive the selection of bacterial variants that are less fit or more vulnerable to other treatments. For example, a phage that uses a multidrug eflux pump as its receptor has been successfully employed to counter-select antibiotic- resistant phenotypes [32].

To better understand the types of phenotypes that may emerge under phage-imposed selection, we investigated the evolution of phage resistance in the model strain *Escherichia coli* MG1655 following exposure to the virulent phage T4. Specifically, we used different protocols to select for genetic variants resistant to T4 infection and analysed the resulting phenotypes. In *E. coli* MG1655, phage T4 requires both LPS and the major porin OmpC to establish infection [33,34]. Because certain antibiotics also rely on porins to enter bacterial cells, and because LPS is an important component for the proper inclusion of porins into the outer-membrane[15], phage predation targeting these structures may select for variants with altered porin expression or structure. This selection, in turn, could coincidentally influence antibiotic susceptibility by decreasing membrane permeability. To explore this possibility, we designed a study to assess whether different selective regimens, ranging from single exposures to prolonged phage pressure, could promote the emergence of bacterial variants with altered susceptibility to clinically relevant antibiotics, including carbapenems and later generation cephalosporins. We show here that the evolution of mutant resistant to phage T4 led to coincidental evolution of antibiotic resistance.

### Bacteriophages use porins and LPS to infect E. coli MG1655

We used the well-characterized laboratory strain *Escherichia coli* K-12 MG1655 as a model to study how selective pressure from phages might promote the emergence of antibiotic resistance. This strain expresses the two major porins, OmpC and OmpF, and displays a truncated rough lipopolysaccharide (LPS) structure lacking the O-antigen due to inactivation of the *wbbL* gene by an insertion sequence [35]. This strain carries no known active antibiotic resistance determinant and is not described to display any virulence factor [36].

The main phage used in this study was the strictly virulent phage T4 [37], a member of the *Tevenvirinae* subfamily (genus *Tequatrovirus*), which efficiently infects *E. coli* MG1655, nearly eliminating the bacterial population within 90 minutes when introduced at a multiplicity of infection (MOI) of 0.1 (Figure 1A). This confirms that under our experimental conditions, phage T4 imposes a strong selective pressure on the bacterial population. We then confirmed that our strain of phage T4 primarily uses the porin OmpC as its receptor and the core LPS side chain as a co-receptor to infect *E. coli* MG1655 as it was described earlier [34]. Deletion mutants lacking *the porin* OmpC (Δ*ompC*) or displaying a deep rough LPS phenotype [38] (*ΔgalU*) showed that both OmpC and core-LPS are required for phage T4 to fully form plaques on *E. coli* MG1655 lawns (figure 1B). The single mutant Δ*ompC* and Δ*galU* reduced the plaques formation by ≈3 orders of magnitude and no plaques was detected for the double mutant Δ*ompC*Δ*galU*. In contrast, the phage T4 fully formed plaques on Δ*ompF* mutant confirming that the alternative major porin OmpF is not necessary for phage T4 infection (Figure 1B). In liquid growth condition, mutations in either Δ*galU* or *ompC* were sufficient to prevent T4 binding and allowed bacterial cultures to grow in the presence of phage T4 (Figure 1C, S1).

**Figure 1:**
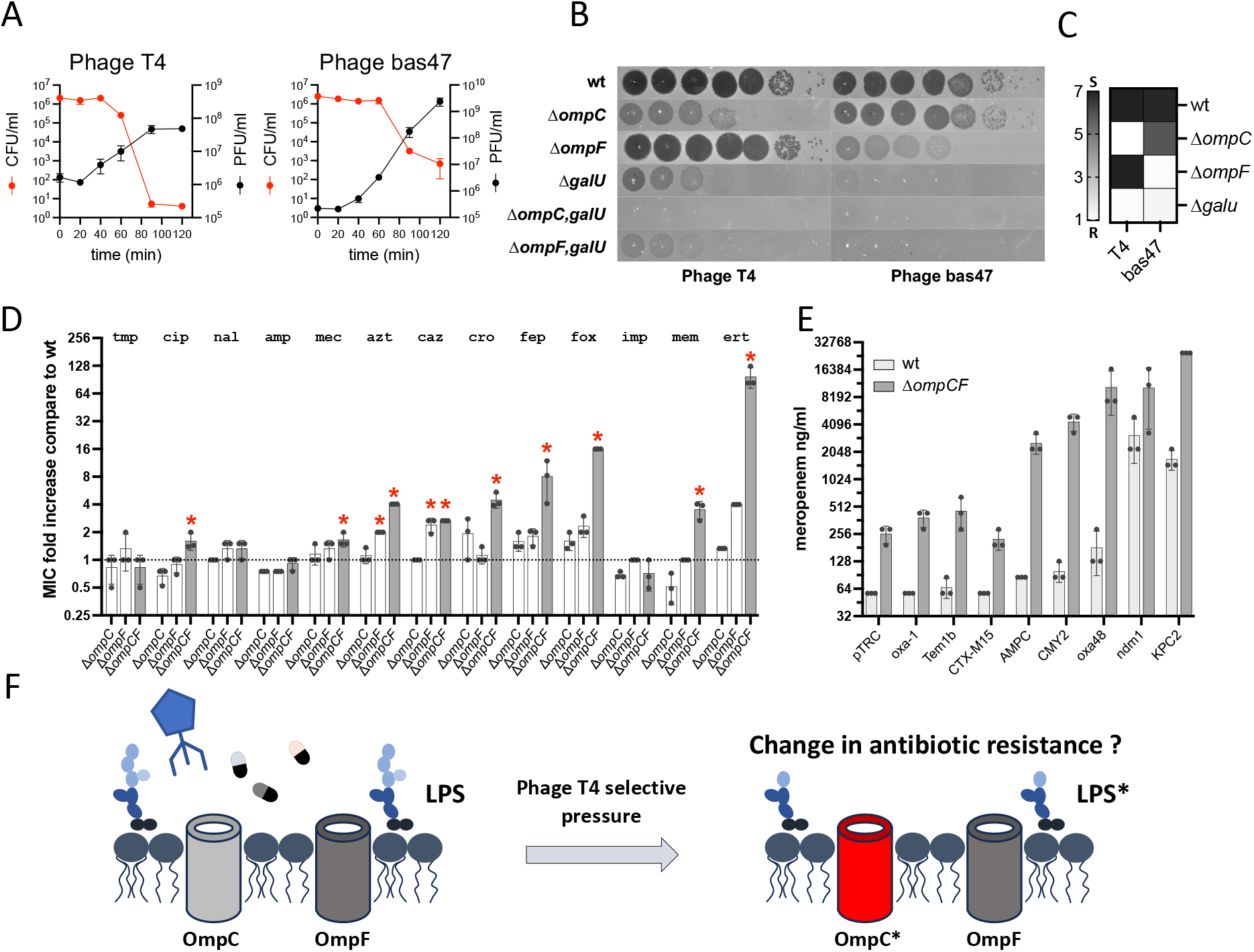
The major porins OmpC and OmpF serve as receptor for bacteriophage and play a role in antibiotic resistance. **(A)** Phages bacteria population dynamics. Each time points represent mean ± sem (n=3). **(B)** Sensitivity of porin and LPS mutants to phage T4 and bas47 measured on semi-solid plaques assay. **(C)** Heat map displaying the phage resistance of the porin and LPS mutants grown in liquid. The resistance R – sensitivity S profile correspond to the ratio at 5h of growth with phage over the growth without phage T4 or bas47 (47) such as ratio of 1 reflect total resistance. **(D)** Antibiotic resistance profile of the porins mutants. The results are display as the fold change of MIC compare to the wt, the dots represent individual replicates value of the E-test reading and the bar the mean ± sd. The red stars illustrate statistical difference (Pvalue <0.05) between wt and Δ*ompCF* tested using multiple comparison one-way ANOVA **(E)** Meropenem resistance profile measured by microdilution of wt or Δ*ompCompF* mutants carrying the pTRC vector expressing beta-lactamases. The dots represent individual replicates of the MIC measures and the bar the ± sd. **(F)** Can phage T4 select for antibiotic resistance ? Here the two major porins are represented together with lipides and LPS structures.

In order to validate our assays and methodology, we additionally included an outlier phage as control. We obtained similar results with phage Bas47 from the Basel phage collection [39], which uses OmpF as a receptor. This phage is related to phage T4 but belongs to a different genus (*Mosigvirus*). In our experimental conditions Bas47 efficiently kills MG1655 (Figure 1A) and relies only on OmpF, not OmpC, to form plaques on *E. coli* MG1655 lawns (Figure 1B). The ability of Bas47 to form plaques was also reduced in a Δ*galU* mutant and was completely inhibited in the double mutant Δ*ompFΔgalU* (Figure 1B). This is either because LPS can act as a co-receptor or because OmpF is not properly displayed in the outer membrane in the deep rough LPS mutant Δ*galU* [40]. Similar to phage T4, a mutation inactivating either OmpF or core-LPS was sufficient to prevent phage attachment (Figure S1A) and allowed bacterial growth in the presence of phage Bas47 in liquid medium (Figure 1C, S1B).

### Impact of *ompC* and *ompF* inactivation on antibiotic resistance

While the role of OmpC and OmpF in antibiotic diffusion is well established [15], the degree of antibiotic resistance induced by their inactivation varies in the literature. To precisely assess the impact of porin inactivation on antibiotic resistance in our experimental set up, we used isogenic *E. coli* deletion mutants lacking *ompC, ompF*, or both genes (Δ*ompC*, Δ*ompF* or Δ*ompCF*). We characterized antibiotic resistance profiles of these mutants by measuring the minimum inhibitory concentrations (MICs) for 13 clinically relevant antibiotics using E-test strips. As shown in Figure 1D and Figure S2, single inactivation of either *ompC* or *ompF* had only a very minor effect on antibiotic MICs, if any. However, the simultaneous inactivation of both porins significantly increased resistance, particularly against cephalosporins (Ceftriaxone, cefoxitin and cefepime) and carbapenems (meropenems and ertapenem). No effect was noted for imipenem.

Clinical studies have reported concerning levels of carbapenem resistance linked to the association between porin loss and expression of certain classes of β-lactamase [17,20,21]. To test this in our experimental setup, we sub-cloned eight β-lactamase (Figure S3A) genes from different families into the same expression vector [41] and measured their impact on meropenem resistance, both in the parental strain and in porins-deficient mutants (Figure 1E). We found that only the two carbapenemase NDM-1 and KPC-2 conferred very high-level meropenem resistance in the parental strain. However, in the Δ*ompCF* double mutant, expression of the cephalosporinase AmpC_CMH4_ and CMY-2, or the oxacillinase OXA-48 led to resistance levels comparable to those observed with NDM-1 or KPC-2. A similar effect, including also CTX-M15, was observed for cefepime (Figure S3B).

β-lactamase inhibitors are essential components of our arsenal to fight against strains producing β- lactamase. We tested also whether porin inactivation could have an effect in the diffusion of Clavulanic acid, Avibactam and Vaborbactam and potentially reduced their activity against β-lactamase in porins deficient mutants. We found that, in our conditions, inactivation of the two majors porins had no significant effect on the activity of β-lactamase inhibitors, and therefore likely not on their diffusion across the outer membrane (Figure S3C).

These findings confirmed that high-level meropenem resistance can be achieved through the loss of both OmpC and OmpF, and further amplified when combined with the expression of β-lactamases from OXA-48 or AmpC family. We next wondered in what measure the selective pressure imposed on porins by certain phages can facilitate the emergence and, or, the spread of antibiotic-resistant bacteria (Figure 1F)?

### Phage T4 selects primarily for mutations in *ompC* and genes related to LPS synthesis

Since phage T4 uses both the porin OmpC and the LPS core to infect *E. coli* MG1655, mutations that disrupt the expression or functionality of these surface structures are likely to be naturally selected under phage lytic pressure. Additionally, mutations in regulatory genes such as *ompR* or *envZ*, which control porin expression may also arise [42,43]. However, inactivation of major porins or disruption of the LPS structure could result in fitness costs, and such mutants may be counter-selected under certain environmental conditions, or during a prolonged cultivation. To explore the range of mutations selected under exposure to phage T4, we performed three independent selection protocols (Figure 2A): i) direct selection of spontaneous mutants by plating bacteria onto phage T4-covered agar plates (T protocol), ii) selection for motile resistant clones by allowing cells to swim through semi-solid medium containing phage T4 (S protocol; Figure S4) and iii) short-term co-evolution using daily bottleneck passages for 4 days in liquid culture (E protocol; Figure S5). We hypothesized that the T protocol, which does not impose a fitness constraint beyond survival, would primarily select for null mutations in *ompC* or LPS synthesis genes. In contrast, the S and E protocols might favour clones with lower fitness costs, such as regulatory mutations or miss-coding mutations, due to additional selection pressures (motility or competitive growth).

**Figure 2.**
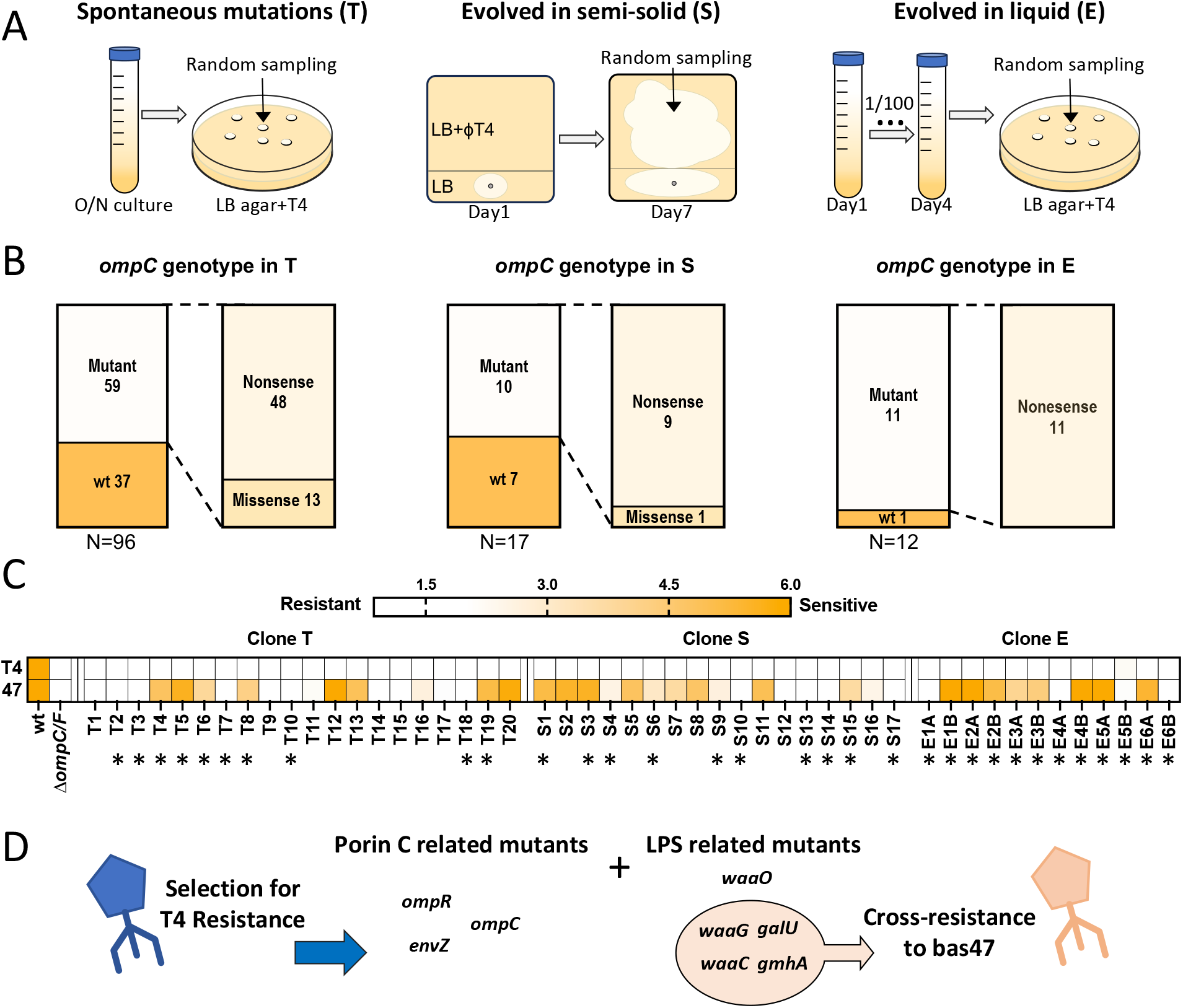
Isolation and characterisation of mutants resistant to bacteriophage T4. **(A)** Representation of the 3 protocols used to select the T4 resistant clone T, S and E. **(B)** Genotype of the *ompC* gene determine by sanger sequencing of the T, S and E clones. **(C)** Heatmap displaying the resistance-sensitivity profile to the phage T4 (T4) and bas47 (47) of the T, S and E clones. The data represent the ratio of the growth of the clones grown with and without phage as such a resistant clone has a value of 1. Clones re-sequenced by NGS are noted by * **(D)** Representation of the main results obtained whole genome sequencing of the T, S and E clones.

We first analysed the *ompC* sequences of 96 T clones, each randomly isolated from an independent culture plated on T4-containing medium. Among these 96 clones, 59 (≈62%) carried mutations in *ompC*, of which 48 (≈85%) were nonsense mutations predicted to abolish OmpC expression (Figure 2B). Next, we analysed *ompC* sequence in 17 independently isolated S clones obtained from swimming plates. Ten of these clones (≈59%) harboured *ompC* mutations, including nine (90%) nonsense mutations (Figure 2B). For the E protocol, we analysed two randomly selected clones per lineage from six independent co-evolution experiments. Of the 12 clones analysed, 11 (≈91%) contained nonsense mutations in *ompC*. Only one lineage featured both clones sharing the same *ompC* mutation, indicating that diverse *ompC* mutations emerged and became fixed independently under constant phage pressure in five out of six co-evolved cultures.

We then selected representative T, S, and E clones based on their *ompC* mutation profiles, including all identified mutation types as well as the parental allele, and confirmed their resistance to phage T4 in liquid culture (Figure 2C, Figure S6). We also included a test of resistance to phage Bas47 to reveal cross resistance phenotypes which could be caused by mutations affecting LPS synthesis pathway or *ompF* expression. (Figure 2C; Figure S6). Interestingly, cross-resistance was mostly found in clones isolated using the protocol T as 60% of clones T, 30 % of clone S and 41% of clones E, also showed cross-resistance to phage Bas47 (Figures 2C; S6; table S1). Notably, some of these cross-resistant clones carried mutations in the *ompC* gene that do not normally impair phage Bas47 infectivity, suggesting the presence of additional mutations outside the *ompC* locus. Conversely, other clones retained the parental *ompC* allele and are thus predicted to harbour mutations in the LPS biosynthesis pathway or in regulatory elements controlling porin expression.

Whole-genome sequencing of a subset of T, S, and E clones (Figure 2D, Table S1) confirmed that cross resistance to phage Bas47 was associated with mutations in genes involved in LPS inner-core synthesis (e.g. *galU, waaG*) but not in side chain or outer core (e.g. *waaJ, waaO*). We found that the T clones, which reflect the selection of spontaneous mutation arising during growth of the bacterial population, contain only mutation in LPS synthesis genes and/or *ompC*. In these clones, miscoding mutations in *ompC* were often associated with additional mutations in LPS synthesis genes. For the S clones we also identified mutations in the EnvZ-OmpR two-component system, which controls porin expression. For the E clones we found that cross resistance was also caused by mutation in the Rcs phosphorelay system, which regulates capsule (colanic acid) synthesis [44]. Taken together, these results highlight that resistance to phage T4 and cross-resistance to phage Bas47 consistently emerge through alterations in surface structures, with porins and LPS pathways representing the dominant genetic targets. These findings demonstrate that multiple, but functionally connected, genetic routes can generate phage resistance, underscoring the central role of envelope composition in shaping bacterial evolutionary responses to phage predation. By combining three independent selection protocols, we show that despite distinct evolutionary trajectories, phage resistance repeatedly converges on mutations affecting porins and LPS biosynthesis, while more complex selection processes additionally favour mutations in regulators of surface antigens.

### Phage-driven selection can potentiate β-lactam resistance

To assess whether phage-driven selection could influence antibiotic resistance, we measured the MIC of meropenem in the 20 T, 17 S, and 12 E mutants using broth microdilution with 1.5-fold serial dilutions (Figure 3A). We selected meropenem as its MIC is particularly affected by porin loss under our conditions (Figure 1D). MICs were quantified using a modified Gompertz fit as described previously [45] (Figure S7A). Overall, most mutants showed only marginal differences in meropenem MIC compared to the wild-type strain. However, since resistance to meropenem typically requires a reduced permeability combined with β-lactamase expression (Figure 1E), we tested whether expressing the *ampC*-type β-lactamase CMY-2 would affect MICs in these genetic backgrounds. In five mutants, CMY-2 expression increased meropenem MIC compared to the ancestral strain (Figure 3A).

**Figure 3:**
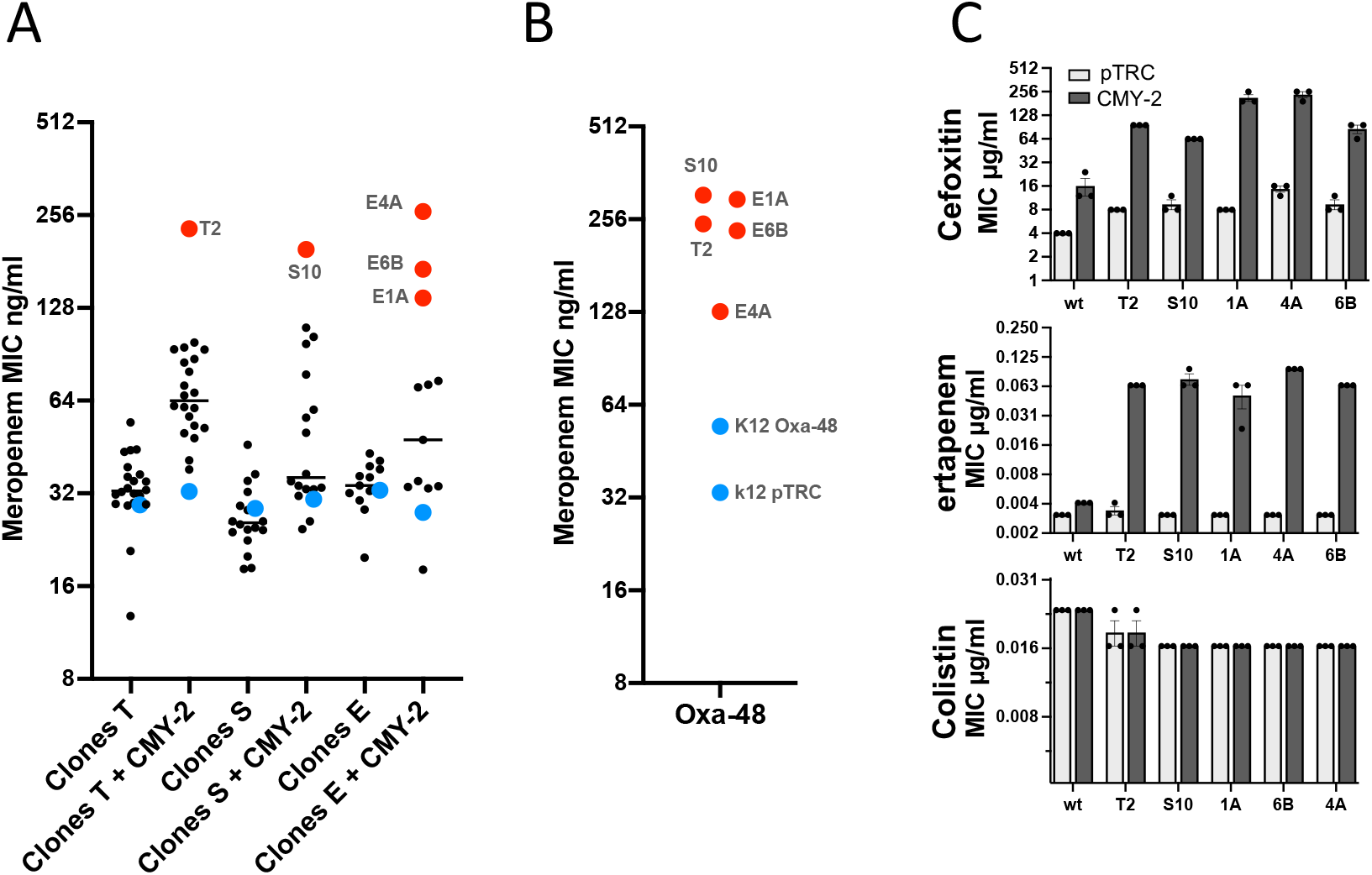
Antibiotic sensitivity of the clone T, S and E. **(A)** Meropenem MIC measured in liquid culture by microdilution for all the clone T, S and E carrying the pTRC vector expressing CMY-2 or not. The Red point correspond to the clone that received further attention. The blue point are the respective baseline wt pTRC and wt CMY-2 **(B)** Meropenem MIC measured in liquid culture by microdilution for all the clone highlighted in (A), carrying the pTRC vector expressing or not OXA-48. **(C)** Ertapenem, cefoxitin and colistin Mic measurement by E-test for the clone selected in (A) with or without the expression of CMY-2

This increase was also observed when these five mutants expressed the OXA-48 β-lactamase (Figure 3B). Further testing using E-test strips showed that these five clones also exhibited altered resistance profiles to cefoxitin and ertapenem when expressing CMY-2. Notably all the 5 clones showed a minor but consistent increased susceptibility to colistin. These results demonstrate that certain phage resistance selected genetic backgrounds, when combined with the expression of β-lactamases, can lead to enhanced resistance to clinically relevant antibiotics.

### Genetic basis of the combined resistance to phage T4, phage Bas47, and β-lactams

All three selection protocols enriched for clones resistant to phage T4, typically through mutations in *ompC* and/or genes involved in LPS synthesis. Some of these mutations also conferred resistance to phage Bas47, suggesting that they simultaneously affected LPS synthesis and porin expression (Figure S7B–D). However, while all mutants with increased antibiotic resistance also displayed cross-resistance to phages, not all phage cross-resistant mutants showed elevated antibiotic resistance. This indicates that only specific combinations of mutations influence both phenotypes. The five clones displaying both increased phage and antibiotic resistance carried nonsense mutations in *ompC* together with mutations in inner-core LPS biosynthesis genes, such as *galU* or *waaG*, resulting in a deep rough LPS phenotype.

To confirm these findings, we constructed mutants with deletions in *waaG* or *waaO*, genes involved in the synthesis of the LPS inner and outer core, respectively, in both the parental strain and a ΔompC background. We then measured their antibiotic resistance with or without CMY-2 or OXA-48 expression (Figure 4). Deletion of *waaG* in a strain expressing OXA-48 was sufficient to raise the meropenem MIC. In the Δ*ompC* background, deletion of *waaG*—but not *waaO*—also increased meropenem MIC when the strain expressed CMY-2 or OXA-48. Since *waaG* mutations were identified in our selection protocol, we further tested whether their inactivation also conferred resistance to other β-lactams. We found that the *waaG* mutation, when combined with *ompC* loss and CMY-2 expression, significantly increased resistance to cefepime, ertapenem, and cefoxitin.

**Figure 4:**
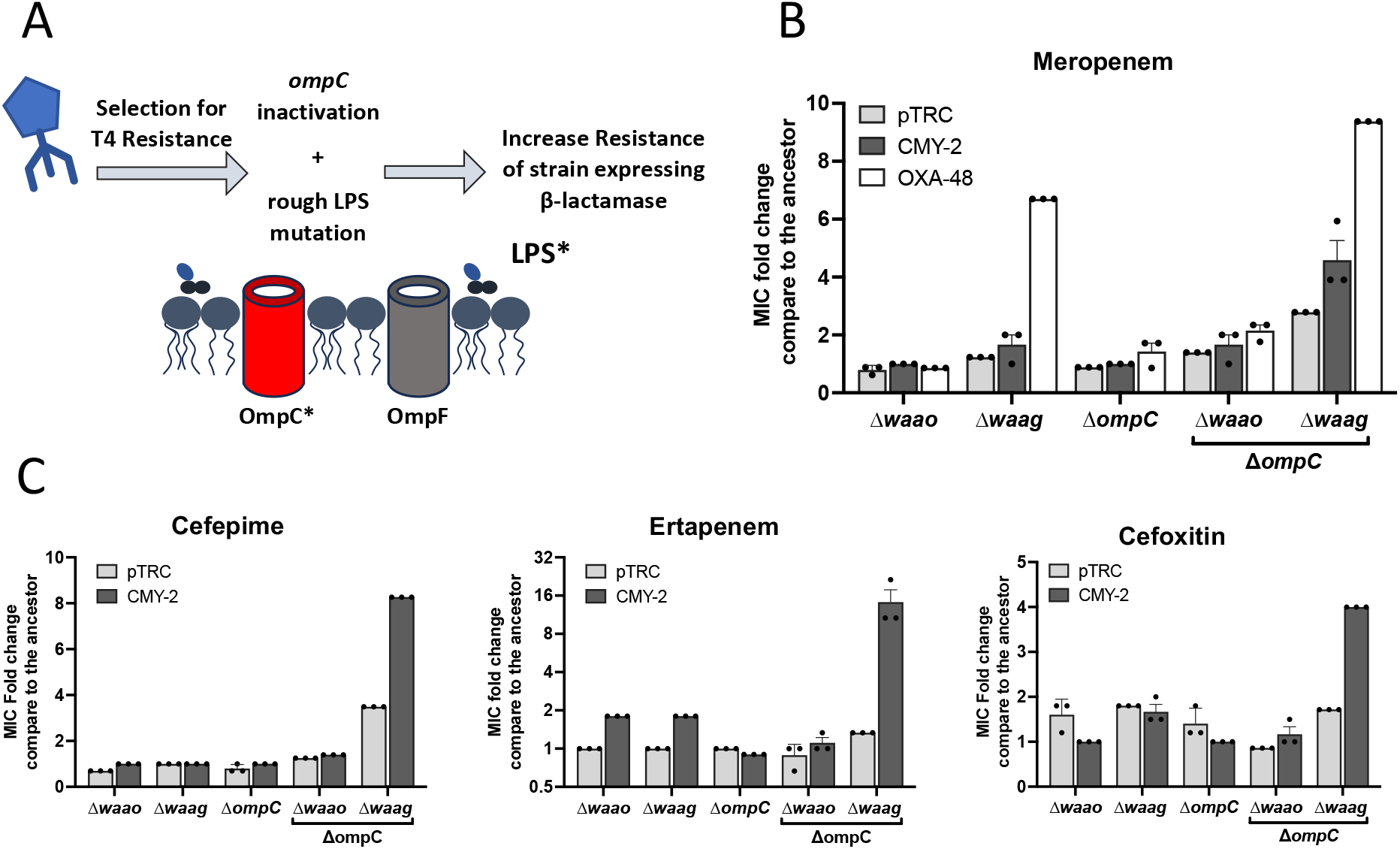
Deep rough LPS mutations associated with OmpC inactivation favour resistance induced by CMY-2 or OXA-48. **(A)** Representation of the mutations synergizing with beta-lactamases found after Phage T4 selection. **(B**,**C)** Fold change in MIC compared to the ancestral strain carrying the pTRC vector expressing or not CMY-2. Mutant strains included total deletions of the ompC gene and/or genes involved in LPS biosynthesis. The wt strain was used as the ancestor for single mutants, while the ΔompC background was used as the ancestor for double mutants.

## Discussion

Phages are recognized as major drivers of bacterial evolution, and their predation imposes a potent selective pressure on surface antigens, which are essential for host recognition and infection. In Gram- negative bacteria, including *E. coli*, phages often rely on outer membrane components such as lipopolysaccharides (LPS) and porins for attachment. These structures also play central roles in environmental adaptation including immune recognition, nutrient uptake and general stress resistance [46,47]. Moreover, LPS and porins regulate antibiotic diffusion inside the bacteria, the LPS through its charge and porins because they serve as entry gates. As such, phage-driven selection on surface receptors likely has pleiotropic consequences, including coincidental shifts in antibiotic resistance profiles.

Here, we demonstrate that selection for resistance to phage T4, which uses both the porin OmpC and the LPS core as receptors, can promote the emergence of *E. coli* genetic backgrounds which favour increased levels of β-lactam resistance, notably carbapenems, when associated with the expression of certain class of β-lactamase. In addition, these selected variants also display cross-resistance to phage Bas47, which exhibits overlapping receptor requirements with phage T4. Whole-genome sequencing revealed that this cross-resistance results from combined mutations in *ompC* and genes involved in the synthesis of the LPS inner core, such as *galU* and *waaG*. Some of these mutations not only prevent the adsorption of both T4 and Bas47 but also likely alter outer membrane permeability, which, when combined with the expression of β-lactamases like CMY-2 or OXA-48, leads to clinically relevant levels of resistance to meropenem, cefepime, and other β-lactams. Both genes are known to disseminate across bacterial species though mobile plasmids.

These results highlight how antagonistic coevolution between phages and bacteria may inadvertently select for antibiotic resistance. The outer membrane of *E. coli* and related *Enterobacterales* is a complex and highly adaptive interface, and its composition can rapidly evolve in response to environmental pressures [47,48]. Notably, LPS architecture is highly variable across enterobacterial species, and even between strains from the same species, owing to the modular biosynthetic pathways [49,50]. This structural plasticity, particularly in the core and O-antigen regions, contributes to phage susceptibility patterns and shapes the evolutionary landscape of receptor usage [51]. Our findings suggest that selective pressure on the LPS core region, often overlooked compared to the O-antigen, may play a critical role in modulating not only phage sensitivity but also antibiotic entry.

Moreover, while porin loss has long been recognized as a mechanism contributing to reduced antibiotic susceptibility, its interplay with phage resistance is still poorly understood. We show that dual inactivation of *ompC* and specific inner-core LPS genes results in a deep rough phenotype that confers resistance to multiple phages while simultaneously impairing the uptake of β-lactams. Importantly, the associated resistance phenotype becomes pronounced only in the presence of β- lactamase expression, consistent with clinical observations that high-level carbapenem resistance often results from the combination of permeability defects and enzymatic degradation [16,20,21].

These findings raise important questions about the potential consequences of therapeutic phage use in clinical settings. While phage therapy holds promise as a targeted alternative to antibiotics, particularly against multidrug-resistant pathogens, our data caution that inappropriate or prolonged exposure to certain phages may unintentionally select for traits that exacerbate antimicrobial resistance. This concern is especially relevant when phages target receptors that also modulate antibiotic permeability, such as porins or LPS components. The long-term evolutionary trade-offs associated with phage resistance, including metabolic costs, changes in virulence, or increased sensitivity to other treatments require careful evaluation on a case-by-case basis. In conclusion, our study illustrates how phage-bacteria interactions can drive the coincidental evolution of clinically relevant phenotypes, including β-lactam resistance. The structural diversity and plasticity of LPS, in particular, emerges as a central element in this evolutionary interplay. Understanding how phages exploit these bacterial features, and how bacteria respond, will be essential for designing effective phage therapies that avoid promoting resistance to conventional antimicrobials

## Materials and Methods

### Bacteria and phage propagation

All the bacterial strains used in this study have been propagated using Lysogenic broth (LB) miller formulation from Difco. Bacterial strains were stocked and inoculated in and from 15% glycerol frozen stocks. LB supplemented with 15g/L bacto-agar from Difco was used as solid medium; soft medium was formed using 7.5g of bacto-agar and swimming medium with 2.8g. When needed for selection kanamycin (100µg/ml), chloramphenicol (30µg/ml) or ampicillin (100µg/ml) was added to the media. Bacteriophages were stocked directly on the LB at 4°C, when needed phage population was amplified on our wt sensitive bacterial strain. All phages and bacteria strain used in this study are listed table S2.

### Gene inactivation using site specific homologous recombination

Deletion mutants of the porin genes and genes involved in LPS synthesis were constructed using site- specific recombination mediated by the λ-Red system, as described by Datsenko and Wanner [52]. The primers listed in table S2 were used to amplify resistance cassettes via PCR using the pkD3 or pKD4 vector as matrices, generating linear DNA fragments containing homology regions for recombination. This linear DNA was introduced into bacteria expressing the λ-Red system carried by the pKD46 vector by electroporation, and recombinant bacteria were selected on the appropriate antibiotic-containing media. Resistance markers were removed using FRT recombination mediated by the expression of the Flp recombinase by the pCP20 vector, as described in the same study.

### Phage and bacteria growth dynamic in liquid

The overnight cultures were diluted 1000-fold in fresh LB medium to a final volume of 5 mL to reach approximately 2-3×10^6^ CFU/mL. 2 × 10^5^ PFU/mL was added to achieve a MOI of 0.1. At each time point, 2 samples of 200 to 500µL were collected. The first sample, to quantify phages, was thoroughly mixed with chloroform at a 10:1 ratio and centrifuged at 14,000 rpm for 5 minutes. The supernatant was recovered and serial dilution were spotted on a soft-agar comprising a lawn of a phage sensitive strain. The second sample, to quantify bacteria, was centrifugated at 14,000 rpm for 5 minutes, the pallet was washed twice with of a solution of MgSO_4_ at 10^−2^ M and the number of CFU was then determined by spotting on LB agar plates.

### Phage sensitivity measure by plaque assay on soft-agar and phage titration

Phages preparations were serially diluted 10-fold, and 5 µL of each dilution was dropped onto soft- agar inoculated from overnight cultures (20-fold dilution) with either the sensitive strain for titration or the tested strain to for sensitivity assessment. The number and appearance of plaque-forming units were recorded after incubation overnight at 37°C.

### Phage adsorption assay

Bacterial cultures me grown at 37°C on flasks at an OD_600_ of 0.4 (≈10^8^CFU/ml) and a control group containing only growth medium were infected with phage at an MOI of 0.001. Infected cultures were kept shaking at 37 and 500µl samples were collected every 2 minutes and mix 10:1 with chloroform. The samples were vortexed thoroughly for 30 seconds and then centrifuged at 14,000 g for 5 minutes at 4°C. The supernatant was serially diluted and spotted on a soft-agar comprising a lawn of a phage sensitive strain.

### Bacterial phage sensitivity profile assayed in microtiter plates

Growth kinetics reflecting the sensitivity to phages were performed on microplates. Overnight cultures were diluted 10000-fold in LB medium containing or not ≈ 10^4^ phage to obtain an MOI of 0.1. After addition of 20µl of the mineral oil plates were incubated at 37°C with orbital shaking in an Infinite® M Nano plate reader and OD_600_ readings were acquired every 5 minutes for 16 hours.

### Subcloning of genes encoding β-lactamases

In order to normalize the expression of the β-lactamases across our conditions, we subcloned them in the pTRC99K vector which derived from the pTRC99A vector and was used already for this purpose to expressed KPC-2 [41,53]. All the constructs are listed table S2, all the new constructs were made using standard restriction/digestion-based cloning of PCR fragments. The PCR fragments including the genes coding β-lactamase were obtained upon amplification of genomic DNA from the NILS collection [54].

### Selection of T4 resistant mutants

The selection of the clones T was done as follow: a total of 96 Independent bacterial cultures were inoculated with up to 10 bacteria and let grow overnight. 5 mL of soft agar was inoculated with ≈10^8^ bacteria and ≈ 10^9^PFU of phage T4. The soft agar was then poured onto LB plate. A Random colony was picked on plates after overnight incubation at 37°C. To assure randomness, plates were marked by a marker spot before incubation, and the closest growing colony from the spot was chosen. The selection of the S clones was done using “dual” swimming squared agar plates consisting on a swimming area without phage and another with phage T4. To create such plates swimming agar was first was poured onto LB agar plate. The LB agar is serving as support to empty two-thirds of the area of the plates from both agar media. Then Swimming agar seeded with 10^9^ PFU/ml of phage T4 was poured in the emptied space. from independent overnight cultures, up to 10 bacteria were seeded in the swimming area without phage. The plates were incubated at 37°C for 3 to 7 days days, and random sampling was performed by collecting bacteria from soft-agar and plating them on LB agar. From these plates a random colony was selected as for the T mutants. The coevolution procedure consisted on starting 6 independent bacterial culture by seeding up to 10 bacteria from an overnight culture in 5ml of LB. upon reaching ≈ 10^8^ CFU/ml the cultures were inoculated with the phage T4 at an MOI ≈ 0.1 and. Daily, for 4 days the bacteria-phage culture was propagated by 100-fold dilution in fresh LB medium. On the final day, after titrating the bacteria, a random two random colonies were selected as for the T mutants.

### Antibiotic resistance assay using microdilution or E-test strip

We used either Microdilution or E-test to assess variation in the MIC of our strains. For microdilution, overnight cultures were diluted 10000-fold in LB medium and dispensed microtiter plate. Serial dilution of the tested antibiotic was added to the well to for a gradient concentration. Microtiter plates were incubated in at 37°C and upon 900 rpm agitation after addition of 10µl mineral oil. After 18 hours of incubation, the OD_600_ was measured using a Synergy H1 plate reader form Biotek. For the E-test strip the procedure recommended by the manufacturer (Biomérieux) was applied in order to obtain a confluent lawn of bacteria. E-test readings were done after 18h incubation.

### Whole genome sequencing and analysis

Bacterial DNA was extracted from the glycerol stocks or overnight culture using the genomic DNA NucleoMag 96 tissue kit from Macherey-Nagel. The whole genome libraries were prepared and indexed using ILMN DNA (M) Tagmentation kit with IDT for Illumina DNA/RNA UD Indexes kit B. The sequencing was performed using Illumina paired-end sequencing 2×150bp on MiniSeq Platform. Illumina reads alignment to the reference genome (NCBI number) and mutation calling was performed using breseq pipeline [55].

### Sanger sequencing

The amplified products or plasmids were checked using Sanger sequencing by Eurofins. Sample purification was performed using Wizard® SV Gel and PCR Clean-Up or Wizard® Plus SV Miniprep DNA purification

## Authors’ contributions

J.LB: data curation, investigation, formal analysis, manuscript review. E.C: Investigation. A.C: Conceptualization, writing—review and editing manuscript, funding acquisition; supervision. A.G.: conceptualization, data curation, formal analysis, funding acquisition, investigation, project administration, supervision, writing—review and editing. All authors read and approved the final manuscript.

## Data availability

All data generated or analysed during this study are included in this published article and its supplementary information files.

## References

1. Clokie MRJ, Millard AD, Letarov AV, Heaphy S. Phages in nature. Bacteriophage. 2011;1: 31–45. doi:10.4161/bact.1.1.14942

2. Chevallereau A, Pons BJ, Van Houte S, Westra ER. Interac(ons between bacterial and phage communities in natural environments. Nat Rev Microbiol. 2022;20: 49–62. doi:10.1038/s41579-021-00602-y

3. Suttle CA. Viruses in the sea. Nature. 2005;437: 356–361. doi:10.1038/nature04160

4. Puxty RJ, Millard AD. Functional ecology of bacteriophages in the environment. Current Opinion in Microbiology. 2023;71: 102245. doi:10.1016/j.mib.2022.102245

5. De Sordi L, Lourenço M, Debarbieux L. “I will survive”: A tale of bacteriophage-bacteria coevolution in the gut. Gut Microbes. 2019;10: 92–99. doi:10.1080/19490976.2018.1474322

6. Suttle CA. Marine viruses — major players in the global ecosystem. Nat Rev Microbiol. 2007;5: 801–812. doi:10.1038/nrmicro1750

7. Koskella B, Brockhurst MA. Bacteria–phage coevolution as a driver of ecological and evolutionary processes in microbial communities. FEMS Microbiol Rev. 2014;38: 916– 931. doi:10.1111/1574-6976.12072

8. Shaer Tamar E, Kishony R. Multistep diversification in spatiotemporal bacterial-phage coevolution. Nat Commun. 2022;13: 7971. doi:10.1038/s41467-022-35351-w

9. Hantke K. Compilation of Escherichia coli K-12 outer membrane phage receptors – their function and some historical remarks. FEMS Microbiology Letters. 2020;367: fnaa013. doi:10.1093/femsle/fnaa013

10. Modulation of host cellular responses by gram-negative bacterial porins. Advances in Protein Chemistry and Structural Biology. Elsevier; 2022. pp. 35–77. doi:10.1016/bs.apcsb.2021.09.004

11. Darby EM, Trampari E, Siasat P, Gaya MS, Alav I, Webber MA, et al. Molecular mechanisms of antibiotic resistance revisited. Nat Rev Microbiol. 2023;21: 280–295. doi:10.1038/s41579-022-00820-y

12. Delcour AH. Outer membrane permeability and antibiotic resistance. Biochimica et Biophysica Acta tiBBA) - Proteins and Proteomics. 2009;1794: 808–816. doi:10.1016/j.bbapap.2008.11.005

13. Baquero F, Marnnez JL, F. Lanza V, Rodríguez-Beltrán J, Galán JC, San Millán A, et al. Evolutionary Pathways and Trajectories in Antibiotic Resistance. Clin Microbiol Rev. 2021;34: e00050–19. doi:10.1128/CMR.00050-19

14. Podolsky SH. The evolving response to antibiotic resistance (1945–2018). Palgrave Commun. 2018;4: 124. doi:10.1057/s41599-018-0181-x

15. Pagès J-M, James CE, Winterhalter M. The porin and the permeating antibiotic: a selective diffusion barrier in Gram-negative bacteria. Nat Rev Microbiol. 2008;6: 893– 903. doi:10.1038/nrmicro1994

16. Nordmann P, Poirel L. Epidemiology and Diagnostics of Carbapenem Resistance in Gram-negative Bacteria. Clinical Infectious Diseases. 2019;69: S521–S528. doi:10.1093/cid/ciz824

17. Poirel L, Potron A, Nordmann P. OXA-48-like carbapenemases: the phantom menace. Journal of Antimicrobial Chemotherapy. 2012;67: 1597–1606. doi:10.1093/jac/dks121

18. Patiño-Navarrete R, Rosinski-Chupin I, Cabanel N, Zongo PD, Héry M, Oueslati S, et al. Specificities and Commonalities of Carbapenemase-Producing Escherichia coli Isolated in France from 2012 to 2015.

19. Oteo J, Delgado-Iribarren A, Vega D, Bautista V, Rodríguez MC, Velasco M, et al. Emergence of imipenem resistance in clinical Escherichia coli during therapy. International Journal of Antimicrobial Agents. 2008;32: 534–537. doi:10.1016/j.ijantimicag.2008.06.012

20. Rosas NC, Wilksch J, Barber J, Li J, Wang Y, Sun Z, et al. The evolutionary mechanism of non-carbapenemase carbapenem-resistant phenotypes in Klebsiella spp. eLife. 2023;12: e83107. doi:10.7554/eLife.83107

21. Bouganim R, Dykman L, Fakeh O, Motro Y, Oren R, Daniel C, et al. The Clinical and Molecular Epidemiology of Noncarbapenemase-Producing Carbapenem-Resistant Enterobacteriaceae: A Case-Case-Control Matched Analysis. Open Forum Infectious Diseases. 2020;7: ofaa299. doi:10.1093/ofid/ofaa299

22. Tamma PD, Goodman KE, Harris AD, Tekle T, Roberts A, Taiwo A, et al. Comparing the Outcomes of Patients With Carbapenemase-Producing and Non-Carbapenemase-Producing Carbapenem-Resistant Enterobacteriaceae Bacteremia. Clinical Infectious Diseases. 2017;64: 257–264. doi:10.1093/cid/ciw741

23. Catacutan DB, Alexander J, Arnold A, Stokes JM. Machine learning in preclinical drug discovery. Nat Chem Biol. 2024;20: 960–973. doi:10.1038/s41589-024-01679-1

24. Czaplewski L, Bax R, Clokie M, Dawson M, Fairhead H, Fischew VA, et al. Alternatives to antibiotics—a pipeline porxolio review. The Lancet Infectious Diseases. 2016;16: 239– 251. doi:10.1016/S1473-3099(15)00466-1

25. Marongiu L, Burkard M, Lauer UM, Hoelzle LE, Venturelli S. Reassessment of Historical Clinical Trials Supports the Effectiveness of Phage Therapy. Clin Microbiol Rev. 2022;35: e00062–22. doi:10.1128/cmr.00062-22

26. Guo Z, Lin H, Ji X, Yan G, Lei L, Han W, et al. Therapeutic applications of lytic phages in human medicine. Microbial Pathogenesis. 2020;142: 104048. doi:10.1016/j.micpath.2020.104048

27. Pires D, Melo L, Vilas Boas D, Sillankorva S, Azeredo J. Phage therapy as an alternative or complementary strategy to prevent and control biofilm-related infections. Current Opinion in Microbiology. 2017;39: 48–56. doi:10.1016/j.mib.2017.09.004

28. Venturini C, Petrovic Fabijan A, Fajardo Lubian A, Barbirz S, Iredell J. Biological foundations of successful bacteriophage therapy. EMBO Mol Med. 2022;14: e12435. doi:10.15252/emmm.202012435

29. Rosas NC, Lithgow T. Targeting bacterial outer-membrane remodelling to impact antimicrobial drug resistance. Trends in Microbiology. 2022;30: 544–552. doi:10.1016/j.tim.2021.11.002

30. Gordillo Altamirano FL, Barr JJ. Unlocking the next generation of phage therapy: the key is in the receptors. Current Opinion in Biotechnology. 2021;68: 115–123. doi:10.1016/j.copbio.2020.10.002

31. Gurney J, Brown SP, Kaltz O, Hochberg ME. Steering Phages to Combat Bacterial Pathogens. Trends in Microbiology. 2020;28: 85–94. doi:10.1016/j.tim.2019.10.007

32. Tarasenko A, Papudeshi BN, Grigson SR, Mallawaarachchi V, Hutton ALK, Warner MS, et al. Reprogramming resistance: phage-antibiotic synergy targets efflux systems in ESKAPEE pathogens. Rodrigues M, editor. mBio. 2025; e01822-25. doi:10.1128/mbio.01822-25

33. Yu F, Mizushima S. Roles of lipopolysaccharide and outer membrane protein OmpC of Escherichia coli K-12 in the receptor function for bacteriophage T4. J Bacteriol. 1982;151: 718–722. doi:10.1128/jb.151.2.718-722.1982

34. Suga A, Kawaguchi M, Yonesaki T, Otsuka Y. Manipulating Interactions between T4 Phage Long Tail Fibers and Escherichia coli Receptors. Johnson KN, editor. Appl Environ Microbiol. 2021;87: e00423–21. doi:10.1128/AEM.00423-21

35. Qin J, Hong Y, Morona R, Totsika M. O antigen biogenesis sensitises Escherichia coli K-12 to bile salts, providing a plausible explanation for its evolutionary loss. Hughes D, editor. PLoS Genet. 2023;19: e1010996. doi:10.1371/journal.pgen.1010996

36. Blattner FR, Plunkett G, Bloch CA, Perna NT, Burland V, Riley M, et al. The Complete Genome Sequence of Escherichia coli K-12. Science. 1997;277: 1453–1462. doi:10.1126/science.277.5331.1453

37. Miller ES, Kutter E, Mosig G, Arisaka F, Kunisawa T, Rüger W. Bacteriophage T4 Genome. Microbiol Mol Biol Rev. 2003;67: 86–156. doi:10.1128/MMBR.67.1.86-156.2003

38. Genevaux P, Bauda P, DuBow MS, Oudega B. Identification of Tn 10 insertions in the rfaG, rfaP, and galU genes involved in lipopolysaccharide core biosynthesis that affect Escherichia coli adhesion. Archives of Microbiology. 1999;172: 1–8. doi:10.1007/s002030050732

39. Maffei E, Shaidullina A, Burkolter M, Heyer Y, Estermann F, Druelle V, et al. Systematic exploration of Escherichia coli phage–host interactions with the BASEL phage collection. Barr J, editor. PLoS Biol. 2021;19: e3001424. doi:10.1371/journal.pbio.3001424

40. Ried G, Hindennach I, Henning U. Role of lipopolysaccharide in assembly of Escherichia coli outer membrane proteins OmpA, OmpC, and OmpF. J Bacteriol. 1990;172: 6048– 6053. doi:10.1128/jb.172.10.6048-6053.1990

41. Amann E, Ochs B, Abel K-J. Tightly regulated tac promoter vectors useful for the expression of unfused and fused proteins in Escherichia coli. Gene. 1988;69: 301–315. doi:10.1016/0378-1119(88)90440-4

42. Slauch JM, Garrett S, Jackson DE, Silhavy TJ. EnvZ functions through OmpR to control porin gene expression in Escherichia coli K-12. J Bacteriol. 1988;170: 439–441. doi:10.1128/jb.170.1.439-441.1988

43. Cowley LA, Low AS, Pickard D, Boinett CJ, Dallman TJ, Day M, et al. Transposon Insertion Sequencing Elucidates Novel Gene Involvement in Susceptibility and Resistance to Phages T4 and T7 in Escherichia coli O157. Fraser CM, Collier RJ, editors. mBio. 2018;9: e00705–18. doi:10.1128/mBio.00705-18

44. Ebel W, Trempy JE. Escherichia coli RcsA, a Positive Activator of Colanic Acid Capsular Polysaccharide Synthesis, Functions To Activate Its Own Expression. J Bacteriol. 1999;181: 577–584. doi:10.1128/JB.181.2.577-584.1999

45. Lambert RJW, Pearson J. Susceptibility testing: accurate and reproducible minimum inhibitory concentration (MIC) and non-inhibitory concentration (NIC) values. J Appl Microbiol. 2000;88: 784–790. doi:10.1046/j.1365-2672.2000.01017.x

46. Simpson BW, Trent MS. Pushing the envelope: LPS modifications and their consequences. Nat Rev Microbiol. 2019;17: 403–416. doi:10.1038/s41579-019-0201-x

47. Lin J, Huang S, Zhang Q. Outer membrane proteins: key players for bacterial adaptation in host niches. Microbes and Infection. 2002;4: 325–331. doi:10.1016/S1286-4579(02)01545-9

48. May KL, Grabowicz M. The bacterial outer membrane is an evolving antibiotic barrier. Proc Natl Acad Sci USA. 2018;115: 8852–8854. doi:10.1073/pnas.1812779115

49. Romeyer Dherbey J, Parab L, Gallie J, Bertels F. Stepwise Evolution of E. coli C and FX174 Reveals Unexpected Lipopolysaccharide (LPS) Diversity. Stern A, editor. Molecular Biology and Evolution. 2023;40: msad154. doi:10.1093/molbev/msad154

50. Leclercq SO, Branger M, Smith DGE, Germon P. Lipopolysaccharide core type diversity in the Escherichia coli species in association with phylogeny, virulence gene repertoire and distribution of type VI secretion systems. Microbial Genomics. 2021;7. doi:10.1099/mgen.0.000652

51. Michel A, Clermont O, Denamur E, Tenaillon O. Bacteriophage PhiX174’s Ecological Niche and the Flexibility of Its Escherichia coli Lipopolysaccharide Receptor. Appl Environ Microbiol. 2010;76: 7310–7313. doi:10.1128/AEM.02721-09

52. Datsenko KA, Wanner BL. One-step inactivation of chromosomal genes in Escherichia coli K-12 using PCR products. Proc Natl Acad Sci U S A. 2000;97: 6640–6645. doi:10.1073/pnas.120163297

53. Compain F, Arthur M. Impaired Inhibition by Avibactam and Resistance to the Ceazidime-Avibactam Combination Due to the D179 Y Substitution in the KPC-2 β-Lactamase. Antimicrob Agents Chemother. 2017;61: e00451–17. doi:10.1128/AAC.00451-17

54. Bleibtreu A, Clermont O, Darlu P, Glodt J, Branger C, Picard B, et al. The rpoS Gene Is Predominantly Inactivated during Laboratory Storage and Undergoes Source-Sink Evolution in Escherichia coli Species. Journal of Bacteriology. 2014;196: 4276–4284. doi:10.1128/JB.01972-14

55. Barrick JE, Colburn G, Deatherage DE, Traverse CC, Strand MD, Borges JJ, et al. Identifying structural variation in haploid microbial genomes from short-read resequencing data using breseq. BMC Genomics. 2014;15: 1039. doi:10.1186/1471-2164-15-1039

