## supplementary figures for "Selection of Escherichia coli porin and LPS mutants under exposure to phage T4 facilitates the emergence of β-lactam resistance"

### Figure S1

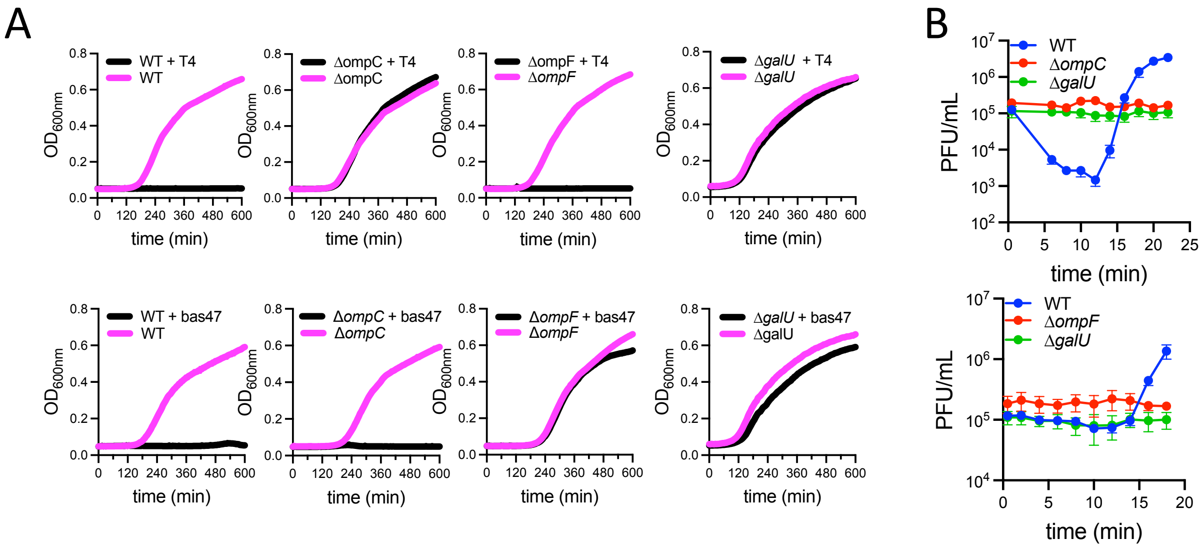

**Figure S1: Resistance-sensitivity of the porins and LPS mutants. (A) )** Representative growth kinetics of the porin and LPS mutants in LB with (black) and without (magenta) the corresponding phage. **(B)** Phages T4 and 47 adsorptions assays against porins and LPS mutant. Each time points represent mean ± sem (n=3).

### Figure S2

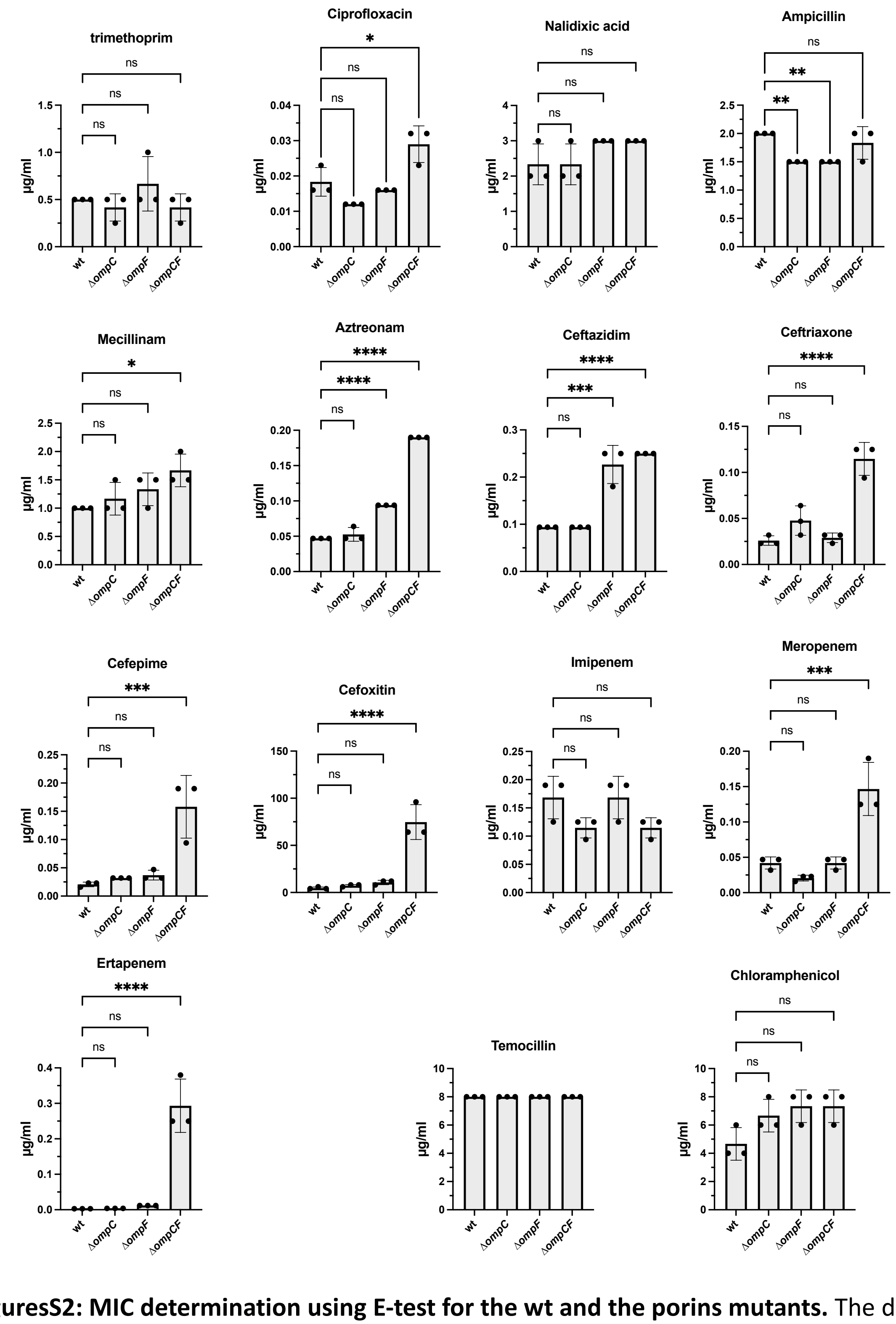

**FiguresS2: MIC determination using E-test for the wt and the porins mutants.** The dots represent individual replicates and the bar the mean ± sd. Statistical significance was tested using multiple comparison one-way ANOVA comparing mean values to mean of the wt. Pvalue * < 0.05 **<0.005 ***<0.0005 ****<0.0001

### Figure S3

A

| Enzyme name | Family | Known feature |
|---|---|---|
| Tem1b | Class A | Broad spectrum |
| CTX-M15 | Class A | ESBL |
| KPC2 | Class A | Carbapenemase |
| NDM-1 | Class B | Carbapenemase |
| $AmpC_{CMH-4}$ | Class C | Cephalosporinase |
| CMY-2 | Class C | Cephalosporinase |
| Oxa-1 | Class D | Broad spectrum |
| Oxa-48 | Class D | Carbapenamase |

B

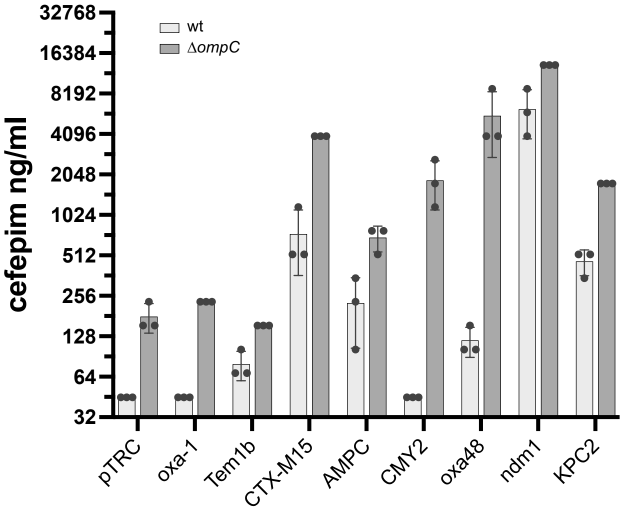

C

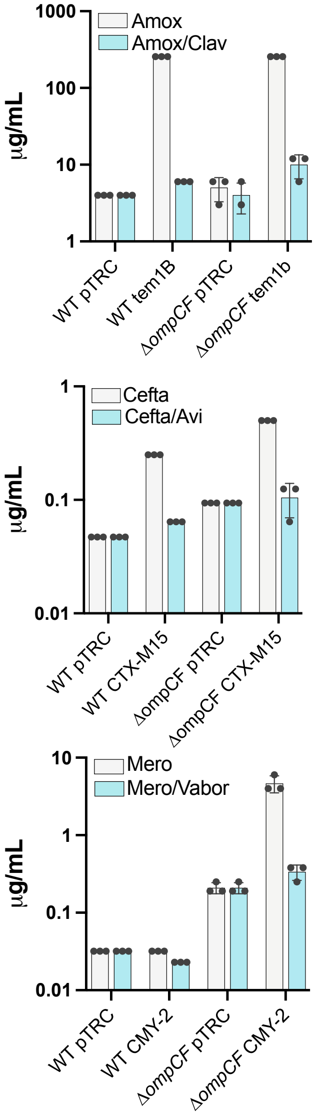

**Figure S3: Impact of porin mutants on beta-lactamase induced beta-lactam resistance and beta-lactamase inhibitor action (A)** List of the beta-lactamase genes used in the study. **(B)** Cefepime resistance profile measured by microdilution of wt or Δ*ompCompF* mutants expressing beta-lactamases. The dots represent individual replicates of the MIC and the bar the ± sd. **(C)** Activity of beta lactamase inhibitor against wt or ΔompCF mutant expressing beta-lactamases Tem1b, CTX-M15, or CMY-2. The dots represent individual replicates value of the MIC measured by E-test and the bar display the mean ± sd.

**Figure S4**

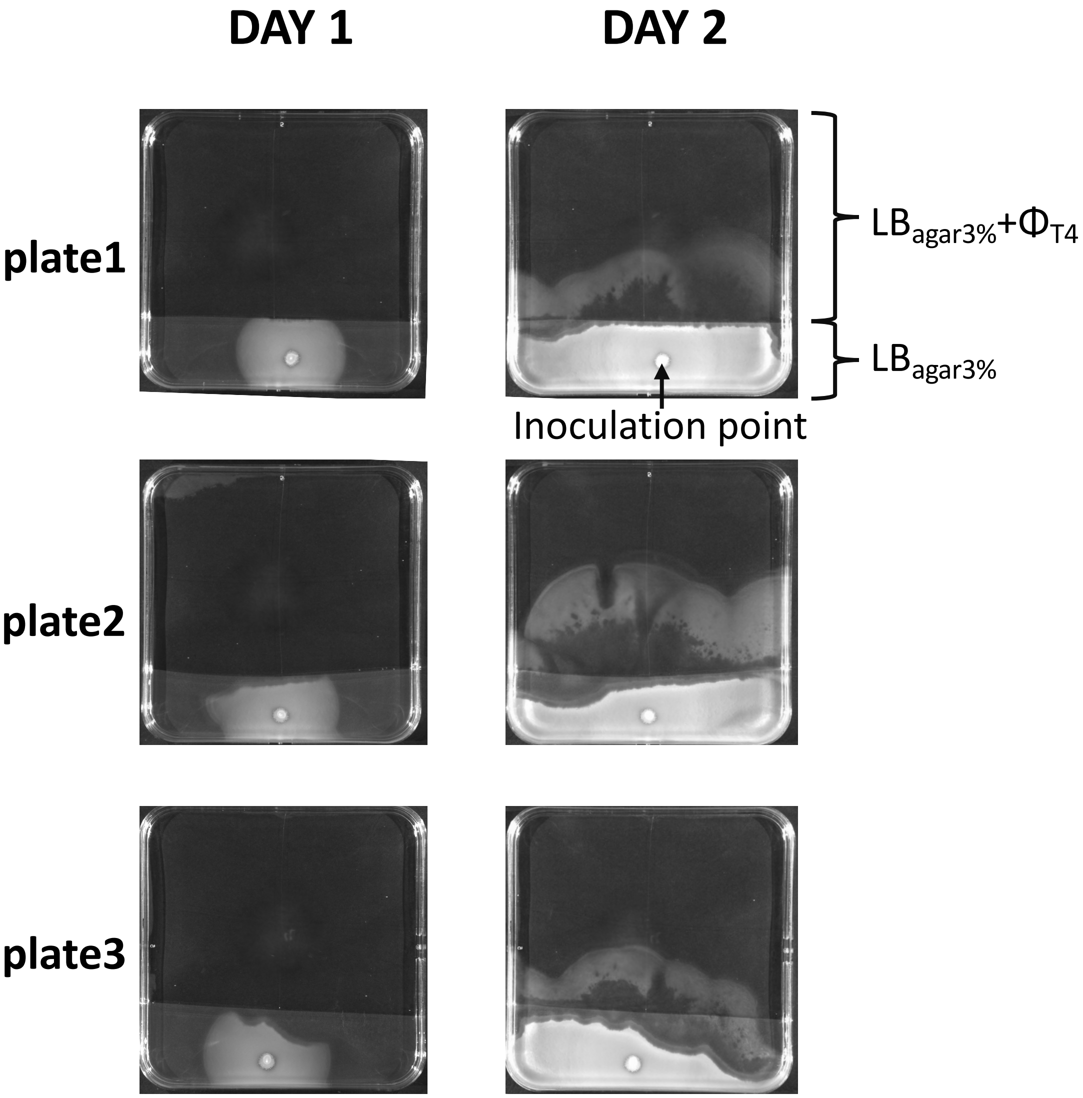

**Figure S4. picture of Dual swimming plates after 24 and 48H of incubation.**

### Figure S5

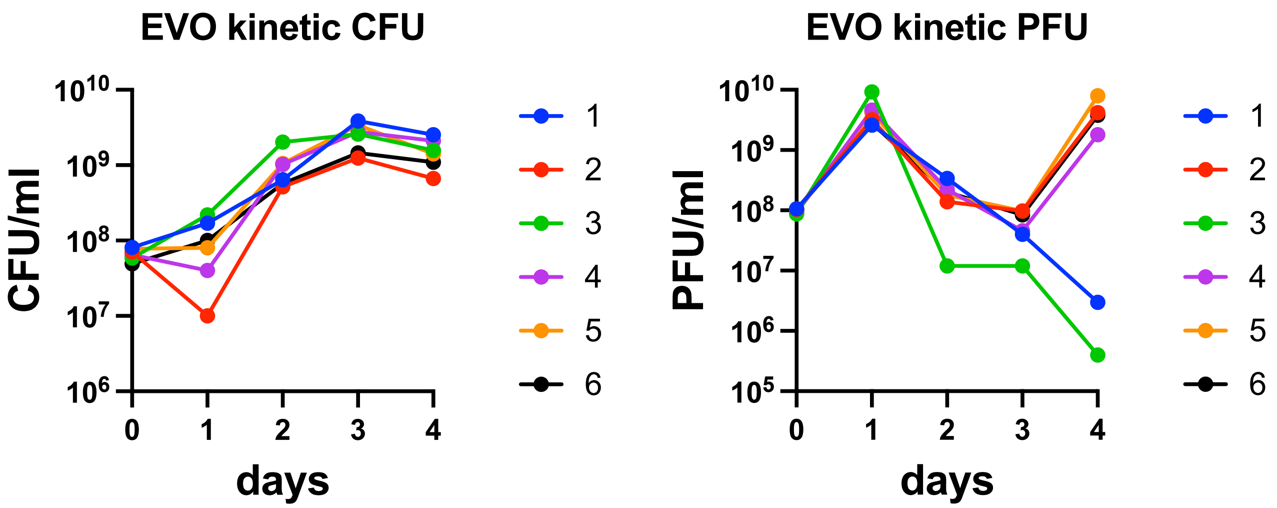

**Figure S5: Phage T4 and Bacteria dynamic across the 4 days of coevolution.** Each colour represent an independent culture. PFU/ml was scored against the ancestor strain.

### Figure S6

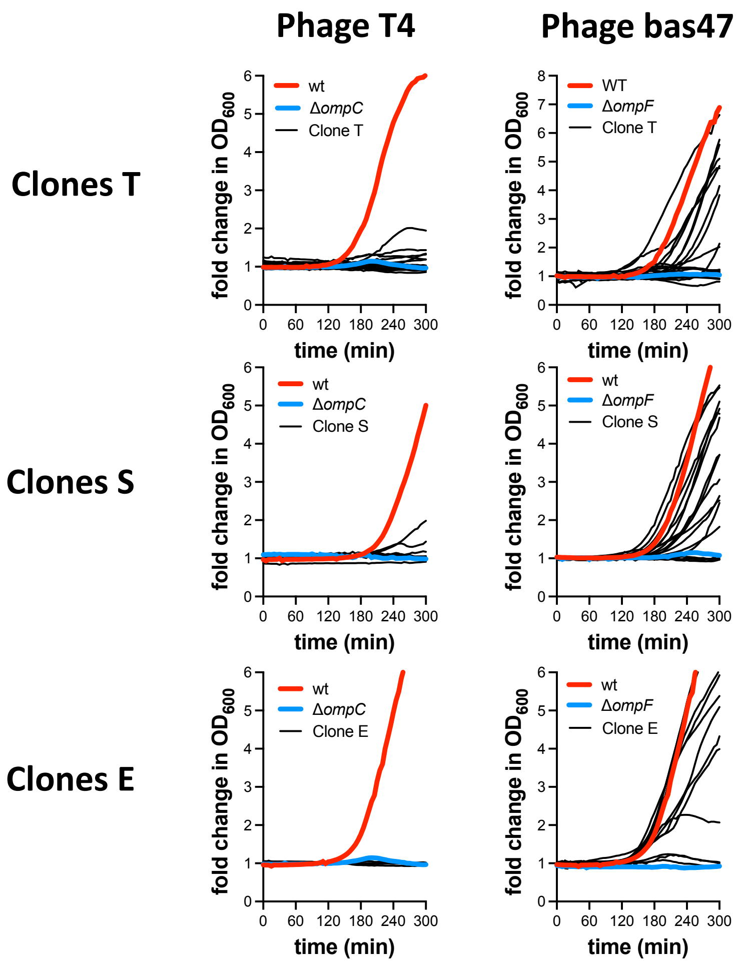

**Figure S6: Fold change over time in $OD_{600}$ of clones T, S, E grown with and without phage T4 or phage 47.**

### Figure S7

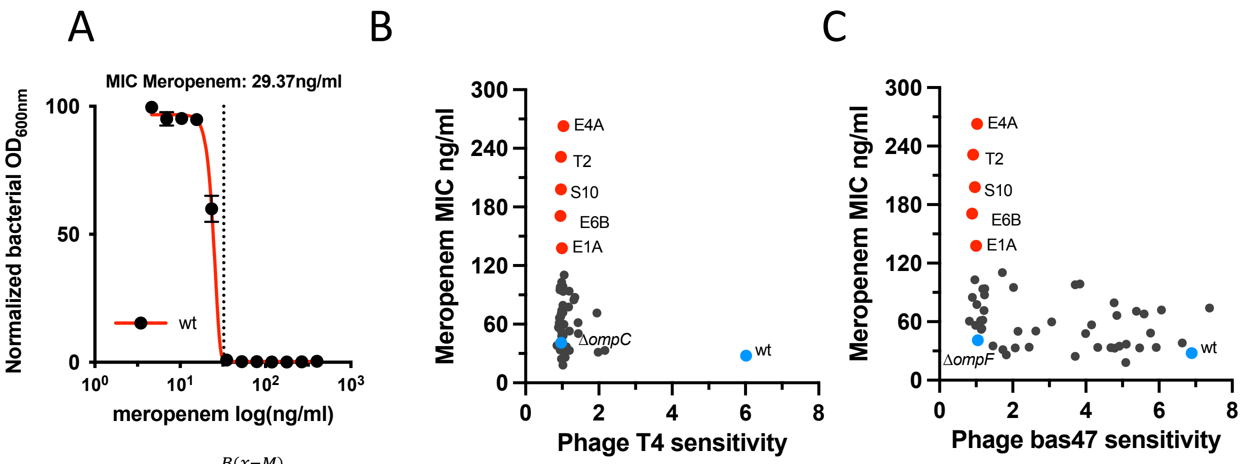

$$y = A + Ce^{-e^{B(x-M)}}$$

$$M = \log \mathrm{MIC} - 1/B$$

**Figure S7: Antibiotic and phge sensitivity of the clone T, S and E. (A)** MIC determination using non-linear fitting of the altered Gompertz function. $A$ is the lower asymptote of $y$, $C$ is the distance between the upper and lower asymptote, $B$ is the slope, and $M$ the log concentration at the inflection point. **(B,C)** Meropenem MIC of the clones T, S and E determined using the function in (A) plotted in function of the T4 (B) or 47 phage resistant profile determined in figure 2C.

| | Isolates | Clones T | *ompC* mutations | LPS mutation | Other mutations | T4 | 47 |
|---|---|---|---|---|---|---|---|
| | 60 | T1 | wt | | | R | R |
| * | 48 | T2* | 6nt del 597-602nt | *waaG* IS862 | | R | R |
| | 56 | T3* | D199N | *waaC* C del 302 | | R | R |
| | 58 | T4* | L292R | *waaO* IS670 | | R | S |
| | 22 | T5* | IS -57nt | | | R | S |
| | 35 | T6* | Q171stop | | | R | S |
| | 57 | T7* | L246R | | | R | R |
| | 16 | T8* | 11nt del 704-714nt | | | R | S |
| | 72 | T9 | wt | | | R | R |
| | 74 | T10* | wt | *waaC* Q160stop | | R | R |
| | 26 | T11 | IS | | | R | R |
| | 10 | T12 | deletion - 1 nucleotide | | | R | S |
| | 53 | T13 | inframe deletion - 81 nucleotides | | | R | S |
| | 84 | T14 | wt | | | R | R |
| | 85 | T15 | wt | | | R | R |
| | 59 | T16 | L296Q | | | R | S |
| | 47 | T17 | inframe deletion - 3 nucleotides | | | R | R |
| | 94 | T18* | wt | *gmhA* +G 484nt | | R | R |
| | 55 | T19* | 93nt del 683-775 | | | R | S |
| | 51 | T20 | 69nt del | | | R | S |

| | clones S | *ompC* mutations | LPS mutation | Other mutations | T4 | 47 |
|---|---|---|---|---|---|---|
| | S1* | 1418nt del ompC- micF-rcsD | *waaO* IS232 | *micF-rcsD* | R | S |
| | S2 | IS -13nt | | | R | S |
| | S3* | wt | *waaO* A del 13nt | | R | S |
| | S4* | Q104stop | waaO IS514nt | | R | S |
| | S5 | IS 35nt | | | R | S |
| | S6* | G185D | *waaO* IS302nt | | R | S |
| | S7 | wt | | | R | S |
| | S8 | wt | | | R | S |
| | S9* | 16nt del 292nt | *waaJ* +T 630 | | R | S |
| * | S10* | IS 8nt | *waaG* IS1003nt | | R | R |
| | S11 | IS 96nt | | | R | S |
| | S12 | IS 11nt | | | R | R |
| | S13* | wt | | *ompR* IS160 | R | R |
| | S14* | wt | *waaj* R112H | *ompR* Q204 stop | R | R |
| | S15* | IS 78nt | *waaO* IS514nt | | R | S |
| | S16 | wt | | | R | S |
| | S17* | wt | *waaF* S17P | *envZ* A239V | R | R |

| | Clone E | *ompC* mutations | LPS mutation | Other mutations | T4 | 47 |
|---|---|---|---|---|---|---|
| * | E1A* | Q54Stop | *galU* IS808nt | | R | R |
| | E1B* | IS -9nt | *waaO* H265Y | | R | S |
| | E2A* | T del 517nt | | *rcsA* T to C -141nt | R | R |
| | E2B* | W273stop | *waaO* IS302nt | | R | S |
| | E3A* | Q171stop | *waaO* IS514nt | | R | S |
| | E3B* | Q171stop | *waaO* IS514nt | | R | S |
| * | E4A* | T del 517nt | *galU* IS221nt | | R | R |
| | E4B* | T to C at -90nt | | | R | S |
| | E5A* | T del 743nt | | | R | S |
| | E5B* | wt | | *rcsD* M186R | R | R |
| | E6A* | Q104stop | waaO IS432 | | R | S |
| * | E6B* | T del 910nt | waaG IS1032 | | R | R |

**Table S1: Comprehensive table to presenting main mutations found using NGS.** The genome of all the clones in red was sequenced by WGS.

| primer name | sequence |
|---|---|
| **Porin deletion** | |
| ompC Fwd | 5' TTGTCTCTTTCTGACGCATTCGTGTTTATGACCGtgtaggctggagctgcttc 3' |
| ompC Rev | 5' CCACGCGGGTAAAGCTTATTTCCATACGTGATTATcatatgaatatcctccttta 3' |
| OmpF Fwd | 5' TAACGCCAGCTTCTAATTTAGCGTCTTCAAGAGGtgtaggctggagctgcttc 3' |
| OmpF Rev | 5' AACATGACGAGGTTCCATTATGGTTACAGAAGGGAAcatatgaatatcctcctta 3' |
| **LPS deletion** | |
| wanner DgalU Fwd | 5'AACACGAACAGTCCAGGAGAATTTAAatgGCTGCCATTAATACGAAAGTCtgcagcattacacgtcttgagcg 3' |
| wanner DgalU Rev | 5' CGTGGATAACACCGATACGGATGttaCTTCTTAATGCCCATCTCTTCTTCAgccatggtccatatgaatatcctcc 3' |
| wanner DwaaG Fwd | 5'CCTTCAGCTGACAGGAATGCACAATTatgATCGTGGCGTTTTGTTTATATtgcagcattacacgtcttgagcg 3' |
| wanner DwaaG Rev | 5' AAAGTGTGGCAAGCGGCTCTTTTAATtcaACCATCTAAACCACCTGTAATaattagccatggtccatatgaatatcctcc 3' |
| wanner DwaaO Fwd | 5'GCTATTTCCCGGAGGAAATAatgCAGCAGGTGTTTTTCCAGGAAACTGAGtgcagcattacacgtcttgagcg 3' |
| wanner DwaaO Rev | 5' TACTTTATAGTTTCCCAGTTttaATGCTTTATCTTTTCAATAAAATAAAAaattagccatggtccatatgaatatcctcc 3' |
| wanner DwaaC Fwd | 5'TACAAGAGGAAGCCTGACGGatgCGGGTTTTGATCGTTAAAACATCGTCGtgcagcattacacgtcttgagcg 3' |
| wanner DwaaC Rev | 5' GATGTTAGCATGTTTTACCTttaTAATGATGATAACTTTTCCAAAACTGCaattagccatggtccatatgaatatcctcc 3 |
| **Beta-lactamase subcloning** | |
| OXA-1 NILS32 Fwd | 5' gcgGAATTCATGAAAAACACAATACATATCAACTTCGC 3' |
| OXA-1 NILS32 Rev | 5' gcgGTCGACTTATAAATTTAGTGTGTTTAGAATGGTGATCGC 3' |
| Tem-1b NILS25 Fwd | 5' gcgGAATTCgcgATGAGTATTCAACATTTCGTGTCGC 3' |
| Tem-1b NILS25 Rev | 5' gcgGTCGACTTACCAATGCTTAATCAGTGAGGC 3' |
| CMY-2 NILS41 Fwd | 5' gcgGAATTCATGATGAAAAAATCGTTATGCTGCGC 3' |
| CMY-2 NILS41 rev | 5' gcgGTCGACTTATTGCAGCTTTTCAAGAATGCGCC 3' |

| Strain name | genotype / feature | source |
|---|---|---|
| WT | E. coli K12 MG1655 | CGSC 6300 |
| ΔompC | Δ*ompC::FRT* | This study |
| ΔompF | Δ*ompF::FRT* | This study |
| ΔompCF | Δ*ompC::FRT* Δ*ompF::FRT* | This study |
| ΔgalU | Δ*galU::FRT* | This study |
| ΔompC,galU | Δ*ompC::FRT* Δ*galU::FRT* | This study |
| ΔompF,galU | Δ*ompF::FRT* Δ*galU::FRT* | This study |
| ΔwaaO | Δ*waaO::FRT* | This study |
| ΔompC,waaO | Δ*waaO::FRT* Δ*ompC::FRT* | This study |
| ΔwaaG | Δ*waaG* | This study |
| ΔwaaG | Δ*waaG::FRT* Δ*ompC::FRT* | This study |
| **DNA source for subcloning** | | |
| NILS 25 | Tem 1b | Bleibtreu et al. https://doi.org/10.1128/jb.01972-14 |
| NILS 32 | oxa-1 | Bleibtreu et al. https://doi.org/10.1128/jb.01972-14 |
| NILS 41 | CMY-2 | Bleibtreu et al. https://doi.org/10.1128/jb.01972-14 |
| **plasmid** | | |
| pTRC99K pTRC | Kanamycin resistance derivatives of pTRC99a | GIFT from Jean-Emmanuelle Hugonnet CRC PARIS |
| pTRC99K ampC | AmpC CMH-4 | GIFT from Jean-Emmanuelle Hugonnet CRC PARIS |
| pTRC99K KPC-2 | KPC-2 | GIFT from Jean-Emmanuelle Hugonnet CRC PARIS |
| pTRC99K NDM-1 | NDM-1 | GIFT from Jean-Emmanuelle Hugonnet CRC PARIS |
| pTRC99K oxa-48 | OXA-48 | GIFT from Jean-Emmanuelle Hugonnet CRC PARIS |
| pTRC99K oxa-1 | OXA-1 | This study |
| pTRC99K tem1B | tem1B | This study |
| pTRC99K CMY-2 | CMY-2 | This study |
| pTRC99K ctx-M15 | CTX-M15 | GIFT from Jean-Emmanuelle Hugonnet CRC PARIS |
| **Cloning vector** | | |
| PKD46 | exo-beta-gam λred | Datsenko and wanner PNAS 2000 |
| PCP20 | FLP recombinase | |
| PKD3 | DNA matrice for chlormaphenicol resistance | |

#### Table S2: Strains and primers used in the study
